# AGS3 antagonizes LGN to balance oriented cell divisions and cell fate choices in mammalian epidermis

**DOI:** 10.1101/2022.05.20.492864

**Authors:** Carlos Patiño Descovich, Kendall J. Lough, Akankshya Jena, Jessica J Wu, Jina Yom, Danielle C. Spitzer, Manuela Uppalapati, Katarzyna M. Kedziora, Scott E. Williams

## Abstract

Oriented cell divisions balance self-renewal and differentiation in stratified epithelia such as the skin epidermis. During peak epidermal stratification, the distribution of division angles among basal keratinocyte progenitors is bimodal, with planar and perpendicular divisions driving symmetric and asymmetric daughter cell fates, respectively. An apically-polarized, evolutionarily-conserved spindle orientation complex that includes the scaffolding protein LGN/Pins/Gpsm2 plays a central role in promoting perpendicular divisions and stratification, but little is known about the molecular regulation of planar divisions. Here, we demonstrate that the LGN paralog, AGS3/Gpsm1, is a novel negative regulator of LGN, and inhibits perpendicular divisions. Static and *ex vivo* live imaging reveal that AGS3 overexpression displaces LGN from the apical cortex and increases planar orientations, while AGS3 loss prolongs cortical LGN localization and leads to a perpendicular orientation bias. Genetic epistasis experiments in double mutants confirm that AGS3 operates through LGN. Finally, clonal lineage tracing shows that LGN and AGS3 promote asymmetric and symmetric fates, respectively, while also influencing differentiation through delamination. Collectively, these studies shed new light into how spindle orientation influences epidermal stratification.

## INTRODUCTION

Asymmetric cell divisions, whereby progenitor cells divide to give rise to daughter cells that adopt different fates, are an important mechanism to promote cellular diversity. Many asymmetric cell divisions rely on proper orientation of the mitotic spindle, which can be influenced by intrinsic and extrinsic cues as well as mechanical forces (Finegan & Bergstralh, 2019; Lechler & Mapelli, 2021; Sunchu & Cabernard, 2020; van Leen, di Pietro, & Bellaiche, 2020; Venkei & Yamashita, 2018; Williams & Fuchs, 2013). Spindle orientation is regulated intrinsically by an evolutionarily conserved ternary complex, which includes the core scaffolding protein LGN (Gpsm2), microtubule binding-protein NuMA/Mud and small G-proteins of the Gαi/o family (Colombo et al., 2003; Du & Macara, 2004; Du, Stukenberg, & Macara, 2001; Schaefer, Shevchenko, & Knoblich, 2000). In *Drosophila*, the LGN ortholog Pins plays a key role in neuroblast stem cells, where it orients the mitotic spindle to promote the unequal inheritance of fate determinants that results in asymmetric daughter cell fates (Bellaiche et al., 2001; Schaefer et al., 2000). The Par3/aPKC/Par6 polarity complex and Insc—among other proteins—facilitate membrane association of the LGN-NuMA-Gαi ternary complex, while downstream, the motor protein dynein mediates pulling forces on astral microtubules to reorient the spindle, (reviewed in Bergstralh, Haack, & St Johnston, 2013; di Pietro, Echard, & Morin, 2016; Morin & Bellaiche, 2011; Tadenev & Tarchini, 2017).

In basal cell progenitors of developing stratified epithelia, we and others have shown that LGN localizes asymmetrically to the apical cortex and promotes perpendicular divisions (Lechler & Fuchs, 2005; Luxenburg, Amalia Pasolli, Williams, & Fuchs, 2011; Williams, Beronja, Pasolli, & Fuchs, 2011; Williams, Ratliff, Postiglione, Knoblich, & Fuchs, 2014). Epidermal loss of LGN, or deletion of NuMA’s microtubule binding domain, leads to elimination of perpendicular divisions, decreased differentiation and impaired barrier function, resulting in neonatal lethality (Seldin, Muroyama, & Lechler, 2016; Williams et al., 2011). These studies highlight the critical importance of perpendicular divisions in establishing proper epidermal architecture. However, only half of mitoses result in perpendicular divisions in wild-type (WT) embryos, with the other half occurring at an orthogonal, planar orientation. LGN is absent from the cortex in planar divisions, and planar divisions occur independently of known spindle orienting proteins, including LGN, NuMA, mInsc, Par3 and Gai3 (Williams et al., 2011; Williams et al., 2014). These observations highlight how little is known about the molecular mechanisms regulating planar divisions, and whether compromising this pathway impacts tissue architecture.

LGN is a member of the AGS (Activator of G-protein signaling) family of proteins, named because they promote G-protein signaling in a receptor-independent manner. In vertebrates, LGN has a closely-related paralog, AGS3 (Gpsm1), and the two proteins share high protein homology and a conserved domain structure, consisting of seven to eight N-terminal tetra-tricopeptide repeats (TPR) and four C-terminal GoLoco motifs—also known as the G-protein regulatory (GPR) region—separated by a flexible linker (Blumer, Cismowski, Sato, & Lanier, 2005; Schiller & Bergstralh, 2021; Wavreil & Yajima, 2020). Biochemical and structural studies predict that AGS3 retains the ability to interact with many of the same binding partners as LGN, including Insc and NuMA (Adhikari & Sprang, 2003; Culurgioni, Alfieri, Pendolino, Laddomada, & Mapelli, 2011; Izaki, Kamakura, Kohjima, & Sumimoto, 2006; Saadaoui et al., 2017; Yuzawa, Kamakura, Iwakiri, Hayase, & Sumimoto, 2011; Zhu, Wen, et al., 2011).

However, there remains sparse evidence that AGS3 possesses spindle orienting activity. An early study showed that AGS3 loss in murine ventricular zone neuronal progenitors increased the proportion of planar divisions, a phenotype that could be mimicked by overexpressing Gαi3 or blocking Gβɣ signaling (Sanada & Tsai, 2005). On the other hand, another study found that LGN, but not AGS3, was expressed in ventricular zone progenitors, and suggested that RNAi targeting of AGS3 had no spindle orientation phenotype (Konno et al., 2008). Most recently, the Morin lab showed that *Gpsm1*^*-/-*^knockout brains have normal spindle orientation (Saadaoui et al., 2017). Overall, while AGS3 does not appear to regulate division orientation in the developing brain, LGN has a well-documented role in promoting planar divisions (Fujita et al., 2020; Konno et al., 2008; Lacomme, Tarchini, Boudreau-Pinsonneault, Monat, & Cayouette, 2016; Mora-Bermudez, Matsuzaki, & Huttner, 2014; Morin, Jaouen, & Durbec, 2007; Saadaoui et al., 2017).

The canonical spindle orientation machinery (including LGN) has long been assumed to operate exclusively during metaphase to orient division prior to chromosome segregation. However, we recently described a novel late-stage spindle orientation process that shares similarity to the post-mitotic reintegration described in Drosophila egg chamber follicular epithelium and murine intestinal crypts (Bergstralh, Lovegrove, & St Johnston, 2015; Cammarota, Finegan, Wilson, Yang, & Bergstralh, 2020; McKinley et al., 2018). We found that a high percentage of epidermal basal cells enter anaphase at oblique angles, which later correct to either planar or perpendicular over the next hour, a process termed “telophase correction” (Lough et al., 2019). While this mechanism is dependent on cell-cell adhesions and their actin scaffolds, it remains unclear whether LGN or other spindle effectors participate in telophase correction or are truly exclusive to early spindle positioning.

Here, we further investigate the role of AGS3 and LGN in regulating oriented cell divisions during epidermal stratification. We find that AGS3 favors planar divisions by antagonizing the ability of LGN to promote and maintain perpendicular divisions. Both knockdown and global knockout of AGS3/Gpsm1 lead to a higher proportion of perpendicular divisions during peak stratification, while overexpression increases planar divisions. AGS3 loss enhances apical LGN localization throughout mitosis, while AGS3 overexpression decreases the efficiency of LGN polarization. Using *ex vivo* live imaging of epidermal explants, we show that LGN not only regulates initial perpendicular spindle orientations during early stages of mitosis, but also plays a maintenance role during telophase, promoting perpendicular reorientation. In support of a model where AGS3 opposes— and acts through—LGN throughout mitosis, we find that AGS3 loss leads to increased perpendicular spindles at anaphase onset and following telophase reorientation, yet AGS3 loss has no effect when LGN is absent. Finally, using mitotic clone genetic lineage tracing in AGS3 and LGN-deficient mice, we show that impairing perpendicular and planar divisions, respectively, has direct effects on asymmetric and symmetric cell fates, while also indirectly impacting the other major route of differentiation in the epidermis, delamination. Together, these data suggest that the two vertebrate Pins orthologs LGN and AGS3 play opposing roles in regulating spindle orientation and differentiation in the developing epidermis.

## RESULTS

### LGN/Gpsm2 and AGS3/Gpsm1 have opposing functions in oriented cell divisions

Proper orientation of the mitotic spindle relies on the evolutionarily conserved ternary protein complex LGN-NuMA-Gai (reviewed in Bergstralh, Dawney, & St Johnston, 2017; Lechler & Mapelli, 2021; Morin & Bellaiche, 2011). During epidermal stratification, this complex localizes to the apical cortex in 50-60% of mitotic basal cells where it promotes perpendicular divisions (Lechler & Fuchs, 2005; Williams et al., 2011; Williams et al., 2014). Previously we have shown that lentiviral knockdown of LGN results in a loss of perpendicular divisions during peak stratification (Lough et al., 2019; Williams et al., 2011). Here, using the spindle midbody protein Survivin to label late-stage (anaphase-telophase) mitotic cells (Fig. 1A,B), we characterized terminal division orientation in embryonic day (E)17.5 back skin epidermis in a variety of loss- and gain-of-function models for LGN (*Gpsm2*) and AGS3 (*Gpsm1*).

**Figure 1.**
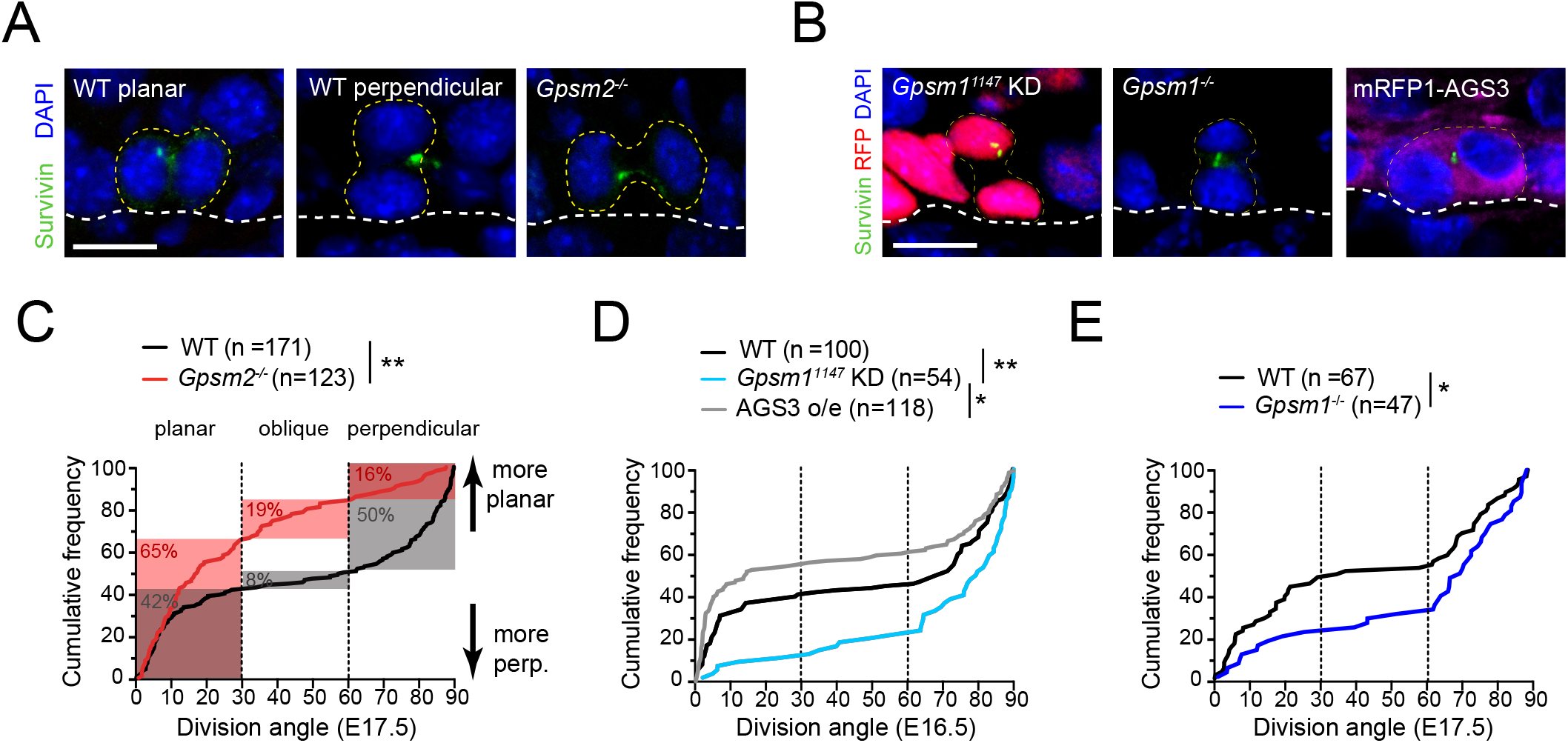
LGN/Gpsm2 and AGS3/Gpsm1 have opposing effects on division orientation. (A) Images from embryonic day (E)17.5 sagittal sections stained for Survivin (green), a late stage mitotic marker that localizes to the spindle midbody during telophase, showing WT planar and perpendicular divisions, and LGN KO (*Gpsm2*^*-/-*^) planar division. (B) Images as in (A) from *Gpsm1*^*1147*^ knockdown, AGS3 KO (*Gpsm1*^*-/-*^), and AGS3-overexpressing telophase cells. (C) Cumulative frequency distribution of terminal division angles from fixed E17.5 sections of WT littermate controls and LGN KO embryos. Planar (0-30°), oblique (30-60°) and perpendicular (60-90°) bins are shown by dashed lines. Shaded areas indicate the proportion of divisions occurring within each bin for WT (black) and *Gpsm2*^*-/-*^KO (red) cells. The upward shift in the LGN KO curve reflects increased planar divisions. (D) Cumulative frequency distributions as in (C). (Left) *Gpsm1*^*1147*^ knockdown (cyan) and uninjected control littermates (black); (middle) AGS3 KO (blue) and WT littermates (black); (right) mRFP1-AGS3 overexpressing (gray) vs uninjected control littermates (black). Scale bars: 10µm. * p<0.05, ** p<0.01 by Kolmogorov-Smirnov test. n values (parentheses) indicate cells from >4 (>5 for overexpression) embryos per genotype. Red (LGN KO), cyan (*Gpsm1*^*1147*^), blue (AGS3 KO) and gray (mRFP1-AGS3 overexpression) color coding will remain consistent throughout figures.

When plotted as a cumulative frequency histogram, wild-type (WT) division angles show a characteristic inverted sigmoid pattern. Most divisions fall within either the planar (0-30°) or perpendicular (60-90°) bins with very few oblique (30-60°) divisions, which accounts for the steep slopes at the beginning and end of the cumulative frequency plot, and the relatively flat middle portion (Fig. 1C, black line). By comparison, the cumulative frequency plot for *Gpsm2*^*-/-*^ knockouts (hereafter, LGN KOs) is shifted upward (Fig. 1C, red line), reflecting an increase in planar and oblique divisions, and a sharp reduction in perpendicular divisions relative to WT controls. The proportions of divisions for each genotype that fall within each orientation bin (planar, oblique, perpendicular) are highlighted by the gray (WT) and red (LGN KO) shaded regions in Fig. 1C. In agreement with our previous analyses of *Gpsm2*^*1617*^ knockdowns (Williams et al., 2011), genetic deletion of *Gpsm2* leads to a significant alteration in division angles, confirming that LGN is necessary for perpendicular divisions.

Despite sharing high homology with its paralog LGN, whether AGS3 possesses spindle orienting activity of its own remains controversial, and two studies in the developing brain have come to different conclusions (Saadaoui et al., 2017; Sanada & Tsai, 2005). Saadaoui and colleagues showed that AGS3 cannot rescue the spindle orientation phenotype caused by LGN loss, while we previously showed that AGS3 loss does not enhance the epidermal thinning phenotype caused by LGN loss (Saadaoui et al., 2017; Williams et al., 2011). Collectively, these data strongly suggest that AGS3 and LGN are not functionally redundant, but leaves unresolved the issue of whether AGS3 plays any role in oriented cell divisions.

To test this, we transduced embryonic epidermis with a lentivirus containing our validated *Gpsm1*^*1147*^ shRNA and a nuclear H2B-mRFP1 reporter (Williams et al., 2011) and evaluated spindle orientation in late-stage mitotic cells at E16.5. Interestingly, upon AGS3 knockdown, we observed a significant downward shift in the cumulative frequency distribution, representing an increase in perpendicular divisions to the detriment of planar divisions (Fig. 1D, compare cyan line to black line). Conversely, overexpression of mRFP1-AGS3 led to a significant increase in planar divisions (compare gray line to black line), phenocopying LGN loss. Like AGS3 knockdowns, we observed an increase in perpendicular divisions in E17.5 germline AGS3 KOs (Fig. 1E, compare blue line to black line). Collectively, these data demonstrate that AGS3 promotes planar divisions during epidermal stratification.

### The LGN/Gpsm2 paralog AGS3/Gpsm1 localizes to the cytoplasm during mitosis

Previous studies across different epithelial tissues have shown that that the subcellular localization of LGN is context-dependent (Ballard et al., 2015; Byrd et al., 2016; Lacomme et al., 2016; Peyre et al., 2011; Williams et al., 2011; Williams et al., 2014). While a recent study has shown that AGS3 is cytoplasmic in mitotic neuronal progenitors (Saadaoui et al., 2017), we took a multi-pronged approach to ascertain the subcellular localization of AGS3 in mitotic epidermal basal cells using immunohistochemistry, lentiviral-mediated expression of tagged proteins, and live-imaging.

A lack of antibodies specific to AGS3 made it challenging to visualize endogenous protein in fixed tissue. However, we were able to circumvent this issue by comparing the staining patterns observed with a validated affinity-purified guinea pig anti-LGN antibody (Williams et al., 2011) and a rabbit pan-LGN/AGS3 antibody (Williams et al., 2014) in WT and LGN (*Gpsm2*) knockout mice. While the latter antibody is commercially available and reported to be LGN specific, another group has suggested it can also recognize AGS3 (Chishiki, Kamakura, Hayase, & Sumimoto, 2017), and similar antibodies raised to the C-terminus of LGN also recognize AGS3 (Konno et al., 2008). Both antibodies detect an apical crescent in WT mitotic E16.5 basal progenitor cells (Fig. 2A, left), consistent with the reported localization of LGN (Lechler & Fuchs, 2005; Lough et al., 2019; Luxenburg et al., 2011; Williams et al., 2011; Williams et al., 2014). However, in mice lacking LGN (Tarchini, Jolicoeur, & Cayouette, 2013), nearly all specific staining was lost for the guinea pig antibody, while cytoplasmic staining remained in mitotic—but not interphase—cells stained with the rabbit antibody (Fig. 2A, right). As further confirmation that the rabbit antibody recognizes AGS3 *in vivo*, we demonstrated that 1) it labels tagged, overexpressed AGS3, and 2) the mitotic cytoplasmic signal we observe in LGN-deficient epidermis disappears when AGS3 is also deleted (Figure 2 – figure supplement 1A,B) Thus, we observed primarily cytoplasmic localization of endogenous AGS3 during mitosis in the developing epidermis.

**Figure 2.**
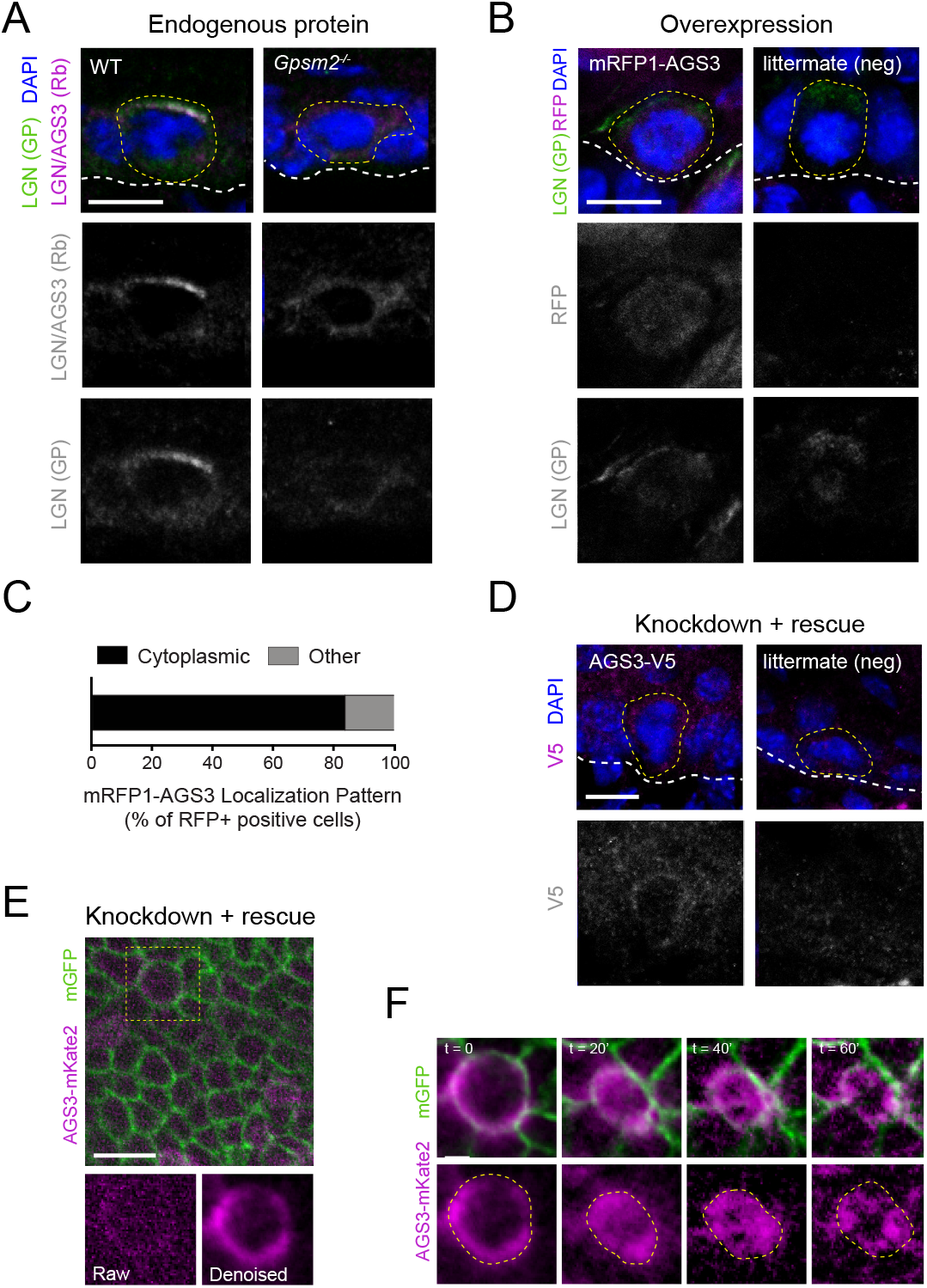
AGS3 localizes to the cytoplasm during mitosis. (A) Immunofluorescent images from E17.5 sagittal sections of wild-type littermate controls (left) and LGN KO (*Gpsm2*^*-/-*^) embryos (right) showing mitotic basal cells stained for guinea pig (GP) anti-LGN (green) and rabbit (Rb) anti-LGN/AGS3 (magenta). Merged images shown at top and single channels below. Cortical signal is lost in LGN KOs while LGN/AGS3 shows cytoplasmic localization in LGN KO cells. (B) E16.5 epidermal basal cell transduced with mRFP1-AGS3 (magenta) and stained for LGN (green). RFP signal shows AGS3 cytoplasmic localization. (C) Fluorescence intensity quantification of whole-cell RFP signal in mRFP1-AGS3 transduced basal cells. Overexpression of mRFP1-AGS3 primarily localizes cytoplasmically. (D) E16.5 epidermal basal cell transduced with AGS3-V5 and stained for V5 (magenta). (E) Image from a movie of a *Krt14*^*Cre*^; *Rosa26*^*mT/mG*^ E16.5 epidermis showing membrane GFP (green), transduced with AGS3-mKate2 (magenta). At bottom, cropped images from yellow dashed box area showing unprocessed (raw) image (left), and denoised/bleach corrected image (right). (F) Denoised stills from movie in (E); t=0 represents metaphase-anaphase transition. Scale bars: 10 µM (A,B,D,F), 50µm (E). * p<0.05 by Mann-Whitney test. Here and in all subsequent figures: dashed white (basement membrane), and dashed yellow line (rough outline of cell borders).

As a second approach, we performed in utero lentiviral injection (Beronja, Livshits, Williams, & Fuchs, 2010) to transduce embryonic epidermis with epitope-tagged AGS3. This method allows high-efficiency (>90%) transduction of surface epithelia using vectors that simultaneously express shRNAs via U6 promoter and fluorescent reporters such as histone H2B-mRFP1 or cDNAs via PGK promoter (Byrd et al., 2016; Dor-On et al., 2017; Lough et al., 2019; Lough et al., 2020; Luxenburg et al., 2011; Luxenburg et al., 2015; Sandoval, Ying, & Beronja, 2021; Williams et al., 2011). First, we overexpressed N-terminal mRFP1-tagged AGS3 (mRFP1-AGS3) in wild-type (WT) embryos and detected its expression in E16.5 dorsal epidermis. This method again revealed cytoplasmic localization in the majority (87%, n = 54) of RFP+ mitotic cells, not seen in uninjected littermates (Fig. 2B,C). Overexpressed mRFP1-AGS3 could be detected through all phases of mitosis (Figure 2 – figure supplement 1C).

Because the addition of a large tag to the N-terminus—where Insc, NuMA and other binding partners interact via the TPR repeats—could interfere with the normal localization or function of AGS3, we also created a lentivirus where AGS3 was tagged with the small V5 epitope at its C-terminus. This construct (AGS3-V5) also contains the validated *Gpsm1*^*1147*^ shRNA (Williams et al., 2011) to knock down endogenous AGS3, and thus accomplishes rescue/replacement rather than overexpression. Immunostaining with a V5 antibody again revealed cytoplasmic localization during mitosis (Fig. 2D).

In order to track the subcellular localization of AGS3 throughout cell division we utilized *ex vivo* live imaging of embryonic epidermal explants (Cetera, Leybova, Joyce, & Devenport, 2018; Lough et al., 2019). We utilized in utero lentiviral transduction to express an shRNA-resistant C-terminal mKate2-tagged AGS3 (AGS3-mKate2) together with the *Gpsm1*^*1147*^ shRNA to knockdown endogenous AGS3. These experiments were performed on a *Krt14*^*Cre*^; *Rosa26*^*mTmG*^ background, which ensured that epidermal cells were membrane-GFP (mG), while *Krt14*-negative tissues (such as dermis and melanocytes) remained membrane-Tomato (mT)+. To eliminate biological artifacts due to phototoxicity, the epidermis could only be imaged at low laser power settings on a Dragonfly spinning disc confocal, which necessitated the implementation of Noise2Void, a deep-learning (DL) image denoising method using self-supervised training (Krull, Vičar, Prakash, Lalit, & Jug, 2020) (Fig 2E). In epidermal basal cells we observed AGS3-mKate2 predominantly in the cytoplasm throughout mitosis (Fig. 2F).

### AGS3/Gpsm1 impacts the cortical localization of LGN/Gpsm2

Biochemical and structural studies have shown that AGS3 can bind to similar interacting partners as LGN, namely Insc, NuMA and Frmpd1 via its N-terminal TPR region, and Gαi proteins via its C-terminal GPR region (Culurgioni et al., 2011; Izaki et al., 2006; Jia et al., 2012; Mauser & Prehoda, 2012; Saadaoui et al., 2017; Yuzawa et al., 2011; Zhu, Wen, et al., 2011). *In vivo* evidence is more nuanced, and elegant chimeric domain-swap experiments between LGN and AGS3 in the developing central nervous system have shown that while the TPR domains of AGS3 can partially substitute for those of LGN in genetic rescue experiments, its linker and GPR domain cannot (Saadaoui et al., 2017). This group also showed that a construct containing the AGS3 TPR region localized to centrosomes—where NuMA is present—while a construct containing the LGN TPR region was cytoplasmic. Moreover, a chimeric protein in which the TPR domains of AGS3 replace those of LGN had a dominant effect, displacing WT LGN from the cell cortex. These studies suggest that LGN and AGS3 might compete for common binding partners.

In mitotic epidermal basal cells, the key spindle orienting cues Insc, Gαi3, and the microtubule-binding protein NuMA all localize to the apical cell cortex with LGN, while the major pool of AGS3 is cytoplasmic, suggesting that any effect AGS3 exerts on oriented cell divisions might be indirect. Collectively, these data led us to hypothesize that AGS3 could displace LGN from the spindle orientation machinery. If correct, altering AGS3 levels should impact the cortical localization of LGN.

To test this, we compared the subcellular localization of LGN following loss or overexpression of AGS3. To do so, we performed lentiviral injection of the *Gpsm1*^*1147*^ shRNA or mRFP1-AGS3, respectively (Fig. 3A). Examination of LGN cortical localization in phospho-histone H3 positive (pHH3+) cells revealed that AGS3 overexpression significantly reduced apical LGN localization and increased the number of cells with unpolarized LGN; conversely, loss of AGS3 resulted in a significant increase of apically recruited LGN (Fig. 3B). These findings are consistent with AGS3 negatively regulating the apical localization of LGN.

**Figure 3.**
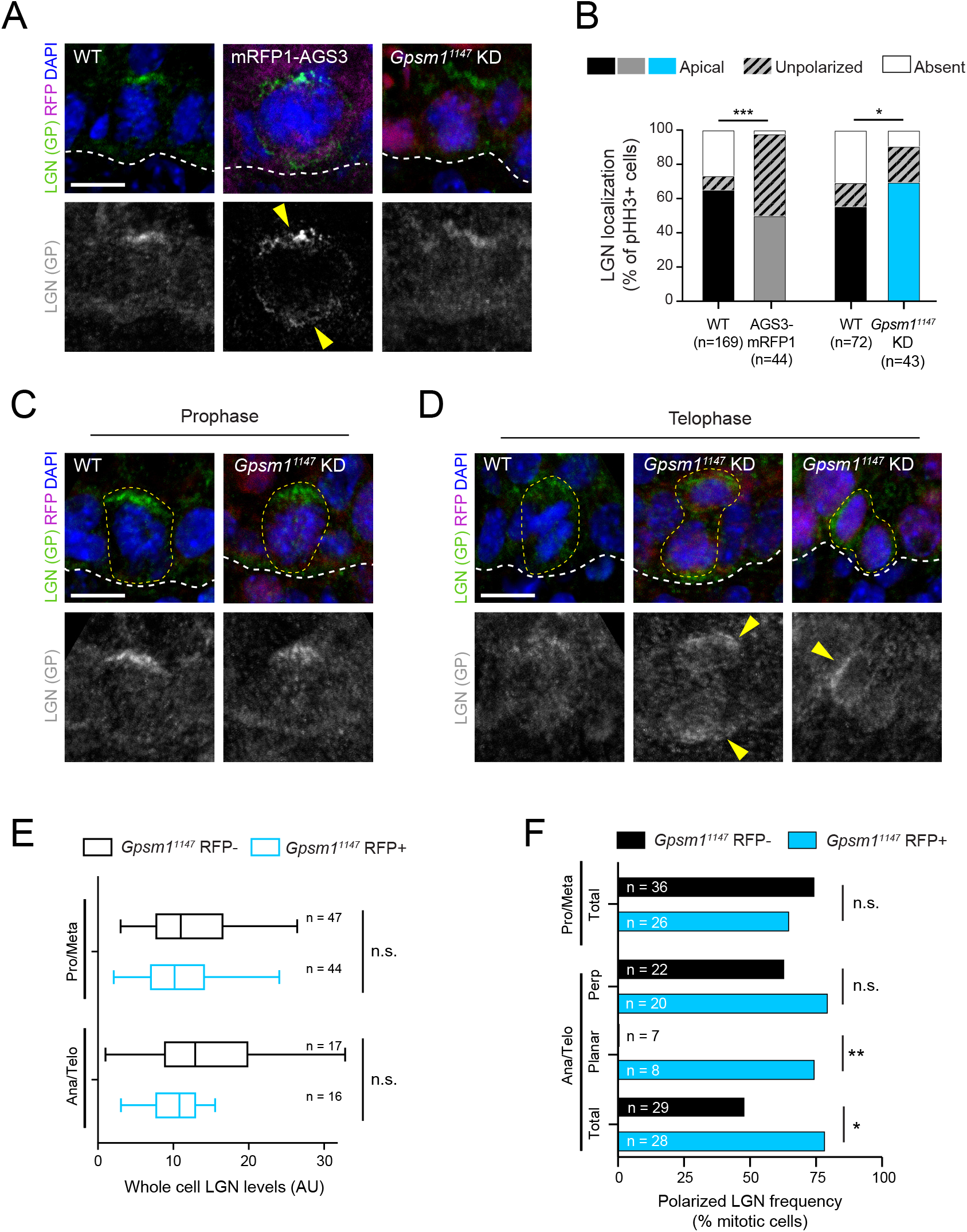
AGS3/Gpsm1 impacts the cortical localization of LGN/Gpsm2. (A) Images of LGN (green) and RFP (magenta) antibody staining from E16.5 sagittal sections of WT (left), mRFP1-AGS3 (center), and *Gpsm1*^*1147*^ knockdown (KD, right) prophase basal cells. Merged images shown at top and single channel LGN (gray) below. Yellow arrows highlight bipolar LGN crescents, an example of an “unpolarized” localization pattern. (B) Quantification of LGN localization patterns, binned by genotype. (C,D) LGN expression (green in merged images and gray below) in prophase (C) or telophase (D) E16.5 epidermal basal cells for indicated genotypes. Yellow arrows indicate atypical LGN expression patterns observed upon *Gpsm1*^*1147*^ KD. (E) Quantification of whole cell LGN levels at indicated mitotic stages. (F) Quantification of frequency that LGN is polarized at indicated mitotic stages for *Gpsm1*^*1147*^ KD (cyan) and WT littermates (black). Ana/telophase cells are split into groups based on division orientation. Scale bars, 10µm; n values indicate cells from >5 (B) or >3 (E,F) embryos/genotype; n.s. = not significant, * p<0.05, *** p<0.001 by Chi-square test (B), Mann-Whitney test (E), or Fisher’s exact test (F).

Next, we sought to determine at which stages of mitosis AGS3 might influence LGN localization. In mosaic tissue transduced with a *Gpsm1*^*1147*^ shRNA, knockdown (RFP+) and wild-type (RFP-) early-stage mitotic cells showed a comparable, high incidence of polarized LGN (Fig. 3C). However, in late-stage mitotic cells, RFP+ cells showed two distinct behaviors compared to RFP-cells: 1) perpendicularly-oriented cells more often showed polarized LGN, sometimes as bipolar crescents, and 2) LGN was often present even in the rare cells with planar orientations (Fig. 3D). Quantification revealed that overall whole-cell levels of LGN were not significantly different between RFP- and RFP+ cells at either stage of mitosis (Fig. 3E). However, the frequency of persistent, polarized LGN was enhanced by AGS3 loss during anaphase/telophase (Fig. 3F).

### LGN/Gpsm2 and AGS3/Gpsm1 play opposing roles during telophase reorientation

Previously, we made the surprising discovery that a significant portion of divisions show oblique (30°-60°) orientations at anaphase onset, but later resolve to planar or perpendicular during a process we call telophase correction (Lough et al., 2019). In WT cells, roughly equal proportions of oblique divisions correct to planar and perpendicular, so any deviation to this ratio suggests an error in reorientation bias. Our finding in fixed tissue that AGS3 loss promoted polarized LGN prompted us to investigate division dynamics in WT, AGS3 KO and LGN KO epidermis using *ex vivo* live imaging (Fig. 4A). To achieve fluorescent labeling of cell membranes in the epidermis, we bred AGS3 and LGN KO mice onto the *Krt14*^*Cre*^; *Rosa26*^*mTmG*^ background, performed confocal imaging on E16.5 epidermal explants at 5-minute intervals, and used Noise2Void to enhance the membrane-GFP (mG) signal.

**Figure 4.**
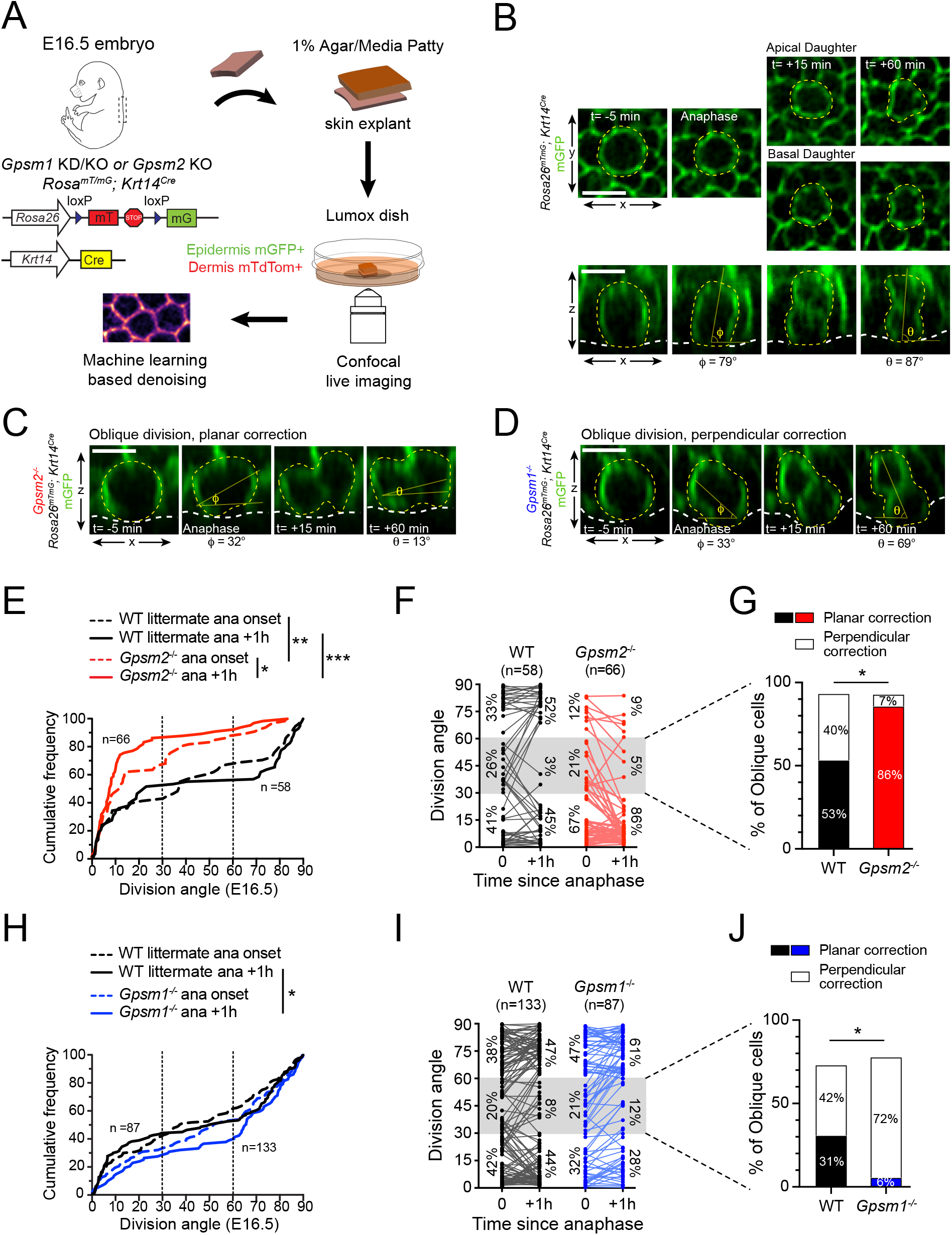
AGS3/Gpsm1 loss biases telophase reorientation toward perpendicular. (A) Schematic for *ex vivo* live imaging of WT, AGS3 KO (*Gpsm1*^*-/-*^) and LGN KO (*Gpsm2*^*-/-*^) embryonic epidermal explants on a *Krt14*^*Cre*^; *Rosa26*^*mT/mG*^ background, where epithelial cell membranes are GFP+. (B) Native en face (top) and z-projections (bottom) movie stills of a WT mitotic cells as it enters anaphase (t= 0), through 1 hour later, depicting a perpendicular division. Division orientation angles are shown below (φ, anaphase onset; θ, +1 h). (C,D) Z-projection movie still from equivalent timepoints showing a LGN KO (C) or AGS3 KO (D) mitosis. See related Figure 4 – figure supplement 1C,D for additional timepoints. (E) Cumulative frequency distribution of division orientation for E17.5 LGN KO embryos at anaphase onset and +1 hr later. (F) Line graphs of individual cell data from (E) depicting orientation at anaphase onset and 1 hr later for LGN KO cells (red) and WT littermate controls (black). Percentages of cells in each orientation bin are shown to the left (t=0) and right (t=+1h) of the data points. (G) Data from (F) depicting behavior of anaphase (t=0) obliques (gray zone in (E)), and frequency of planar vs perpendicular correction; rare cells that remain oblique are not included. (H-I) Similar plots as (E-G) for AGS3 KO cells (blue) and WT littermate controls (black). Scale bars, 10 µm; n values indicate events from 3-4 embryos imaged in 2 technical replicates per genotype; * p<0.05, ** p<0.01, *** p<0.001 by Kolmogorov-Smirnov test (E,H) or Fisher’s exact test (G,J).

An example of a WT basal cell undergoing a perpendicular division, shown in both the native en face (xy) and z-projection (xz) views can be seen in Fig. 4B. Examples of representative LGN KO and AGS3 KO cells are shown in Fig. 4C,D, while additional examples of movie stills in different genotypes can be found in Figure 4 – figure supplement 1A-D. In agreement with our previous studies (Lough et al., 2019)—conducted on a different genetic background with deconvolution post-processing rather than denoising—as a population, WT cells displayed “randomized” division orientation at anaphase onset but corrected to a bimodal/inverse sigmoidal profile by 1h later (Fig. 4E, black lines).

Our previous data showed that LGN knockdown led to an increase in the percentage of cells that enter anaphase with planar spindles (Lough et al., 2019), which was expected because apical LGN promotes perpendicularly-oriented spindles during early mitosis (Williams et al., 2011). While LGN knockdown cells exclusively corrected toward planar, we were unable to make any definitive conclusions about a “maintenance” role for LGN during telophase reorientation because of the small number of cells which entered anaphase at >30°. Here, we address this question by collecting a larger live-imaging dataset of LGN KO epidermal explants, where de-noising also enhanced our ability to detect the mG membrane signal and measure division angles in z-projections with precision.

Compared to WT, LGN KO basal cells showed a strong bias toward planar and oblique orientations at anaphase onset, which was significantly accentuated 1h later (Fig. 4E, red lines). At the individual cell level, WT cells entered anaphase at planar (0°-30°, 41%), oblique (30°-60°, 26%) and perpendicular (60°-90°, 33%) angles in roughly equal proportions (Fig. 4F). By comparison, in LGN KO mutant explants, 67% of cells entered anaphase at planar orientations, confirming that genetic loss of LGN leads to a similar increase in planar-oriented spindles as we observed upon *Gpsm2*^*1617*^ knockdown. Of note, among the obliquely-oriented spindles, correction was equally likely in either direction in WT explants, while in LGN KO explants, >85% (12/14) corrected to planar (Fig. 4G, example shown in Fig. 4C). These data strongly suggest that apical LGN not only directs initial spindle positioning during early mitosis, but also promotes perpendicular reorientation during late mitosis.

Next, we performed the same experiments on a AGS3 KO background to determine whether AGS3 regulates initial spindle positioning, reorientation, or both. Once again, WT littermate controls refined from an evenly-distributed to a bimodal pattern during the anaphase-telophase transition period (Fig. 4H, black lines). At the population level, the distribution of division angles in AGS3 KOs displayed a downward shift—indicating a perpendicular bias compared to WT littermates (Fig. 4H). At the individual cell level, WT obliques were once again similarly likely to correct to planar or perpendicular. However, among AGS3 KO obliques, the vast majority (72%, 13/18) corrected to perpendicular (Fig. 4I,J; example shown in Fig. 4D). These data demonstrate that while LGN influences both initial spindle positioning and reorientation, the effect of AGS3 is most pronounced in telophase correction. Moreover, while LGN promotes perpendicular reorientation, AGS3 promotes planar reorientation.

### Gpsm2 (LGN) is epistatic to Gpsm1 (AGS3)

We have shown that AGS3 overexpression or loss can inhibit or enhance, respectively, the apical localization of LGN, and that AGS3 and LGN have opposing effects on telophase correction during oriented cell divisions. If AGS3 inhibits the activity of LGN, then LGN would act downstream of AGS3 and we would predict that *Gpsm1* loss should not affect the *Gpsm2* mutant phenotype. To test this, we compared spindle orientation phenotypes—in fixed tissue and using *ex vivo* live imaging—caused by dual loss of LGN and AGS3 to loss of LGN alone.

We generated two dual loss-of-function models for AGS3 and LGN. First, we used our in utero lentiviral transduction to generate mosaic *Gpsm1*^*1147*^knockdown on a *Gpsm2*^*-/-*^ null background and compared RFP+ (double loss-of-function) to RFP-(LGN single KO) populations. As an alternative, we interbred the *Gpsm1* and *Gpsm2* lines to create germline double KOs. Cumulative frequency histograms of Survivin+ terminal-stage mitotic cells revealed strong biases toward planar divisions in all groups, with no significant differences between them (Fig. 5A,B). Thus, loss of AGS3 has no effect on the LGN phenotype of increased planar divisions in fixed tissue.

**Figure 5.**
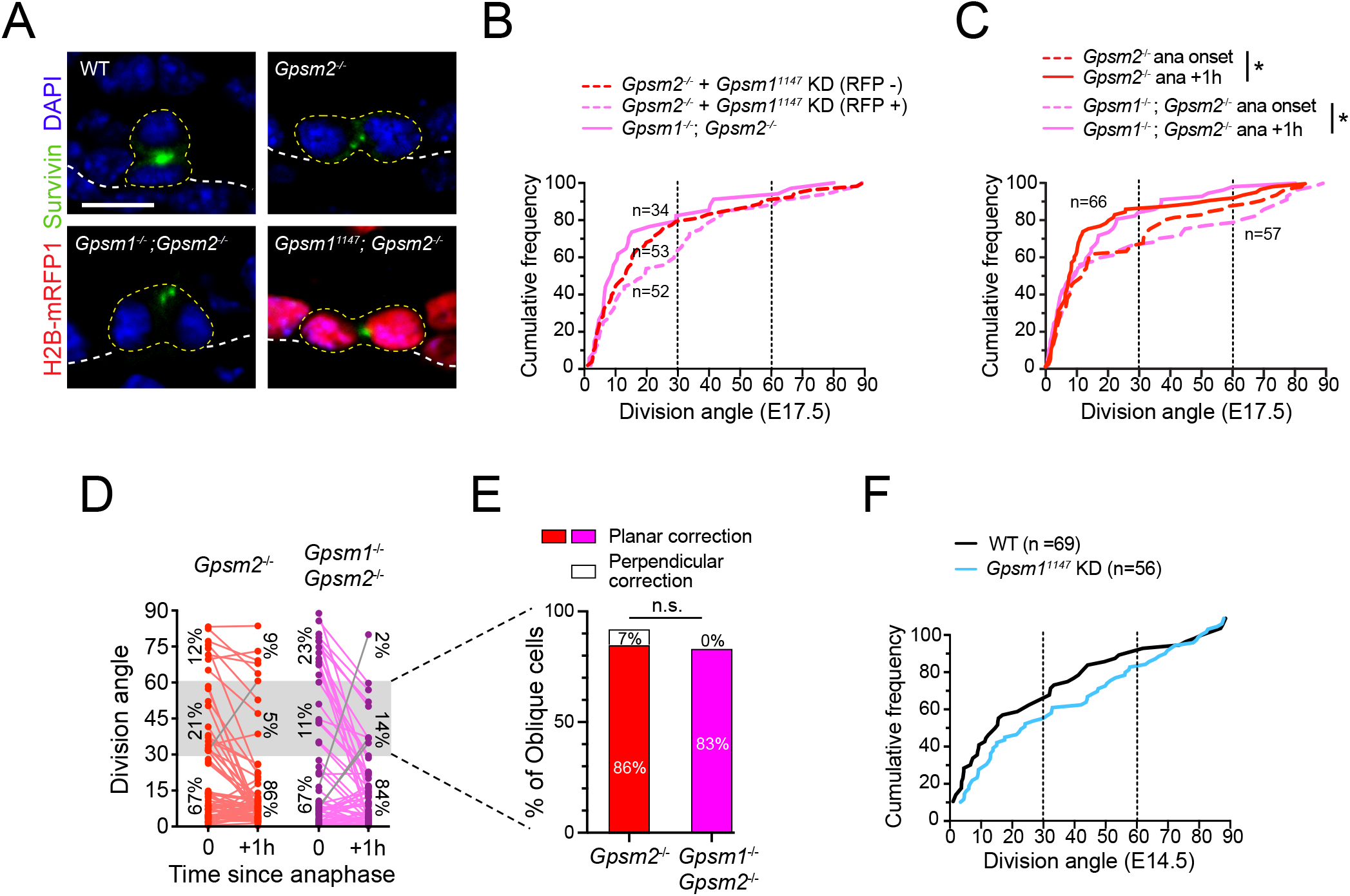
Gpsm2 is epistatic to Gpsm1. (A) Images of telophase cells from E17.5 WT, LGN KO (*Gpsm2*^*-/-*^), AGS3/LGN double knockout (*Gpsm1*^*-/-*^; *Gpsm2*^*-/-*^) and AGS3 KD/ LGN KO (*Gpsm1*^*1147*^; *Gpsm2*^*-/-*^) backskin epidermis, labeled with the late-stage mitotic marker Survivin (green). (B) Cumulative frequency distribution of telophase division angles from fixed sections for indicated genotypes: *Gpsm2*^*-/-*^; *Gpsm1*^*1147*^ H2B-RFP-(LGN KO, red dashed line), *Gpsm2*^*-/-*^; *Gpsm1*^*1147*^ H2B-RFP + (AGS3 KD/LGN KO, magenta dashed line), and *Gpsm1*^*-/-*^; *Gpsm2*^*-/*^(AGS3/LGN double KO, magenta line). (C) Cumulative frequency distribution of division angles at anaphase onset (dashed lines) and 1h later (solid lines) from live imaging experiments of LGN single KOs (red, replotted from Fig. 4E) and AGS3/LGN double mutants (magenta) (D) Line graphs of individual cell data from (C) depicting orientation at anaphase onset and 1 hr later for LGN KO (red) and AGS3/LGN double KO cells (magenta). (E) Data from (D) depicting planar vs perpendicular telophase correction behavior for anaphase obliques (gray bar), showing that 83% of double knockout cells correct to planar, similar to the 86% observed in LGN single KOs. (F) Cumulative frequency distribution of telophase division angles quantified from E14.5 back skin sections for *Gpsm1*^*1147*^ knockdown H2B-RFP+ cells (cyan) and WT uninjected littermate controls (black). Scale bars, 10 µm; n value indicates cells from >3 independent embryos per genotype (B) with at least 2 technical replicates (C-E); n.s. = not significant; * p<0.05, by Kolmogorov-Smirnov test (B,C) or chi-square test (E).

Next, we crossed *Gpsm1* and *Gpsm2* alleles onto the *Krt14*^*Cre*^; *Rosa26*^*mT-mG*^ background to perform *ex vivo* live imaging. Due to complex breeding schemes and the six alleles needed to generate these mice—as well as the small litter sizes obtained from AGS3 KOs—we were not able to obtain LGN single KOs from the same litters. Thus, we compare the double mutants imaged in this experiment to the same LGN KO group shown in Fig. 4C-E. On cumulative frequency histograms, both LGN single KO and AGS3/LGN double KO populations showed a similar significant “left-ward” shift toward increasing planar divisions 1h after anaphase onset, with nearly overlapping curves (Fig. 5C). At the individual cell level, cells were twice as likely to enter anaphase at planar compared to non-planar orientations for both LGN single KOs and AGS3/LGN double KOs, and correction was almost universally planar in both groups (exceptions shown in gray in Fig. 5D). Finally, among anaphase obliques, over 80% in both genotypes corrected to planar (Fig. 5E). These data confirm that *Gpsm2* is epistatic to *Gpsm1* in telophase correction.

Finally, we reasoned that if AGS3 required LGN for its function, then AGS3 loss should have no effect when LGN is dispensable. To test this, we took advantage of the fact that epidermal stratification occurs in both LGN-independent and LGN-dependent phases. In the single-layered epithelium (E12.5 and earlier), divisions are initially planar, become “randomized” by E13.5-E14.5 when stratification commences, and finally adopt their mature “bimodal” distribution by ∼E15.5-E16.5 (Damen et al., 2021; Lechler & Fuchs, 2005; Williams et al., 2014). LGN first shows apical polarization at ∼E15.5, and LGN loss has no effect on spindle orientation prior to this age (Williams et al., 2014). Thus, the initial phases of stratification occur independently of LGN. Interestingly, like LGN, loss of AGS3 in early (E14.5) epithelia caused no significant effect on spindle orientation (Fig. 5F). Collectively, these data show that AGS3 loss has no apparent effect when LGN is absent, and strongly suggest that AGS3 acts through LGN in spindle orientation.

### AGS3/Gpsm1 loss promotes asymmetric cell fates and differentiation

Spindle orientation is frequently, but not always linked to cell fate choices (Williams & Fuchs, 2013). Correlative studies have shown that loss of core apical complex proteins such as Gαi3, Insc, Par3, NuMA and LGN—which reduce perpendicular divisions—also lead to epidermal thinning, thus demonstrating that they impact stratification (Seldin et al., 2016; Williams et al., 2011; Williams et al., 2014). Our *ex vivo* live imaging studies have revealed previously unappreciated plasticity in daughter cell positioning during late stages of mitosis. Thus, it is possible that daughter cells could either differentiate following divisions or alternatively, reintegrate into the basal layer following mitosis. Such behaviors have been recently documented in other epithelia (McKinley et al., 2018; Wilson & Bergstralh, 2017), and dedifferentiation has been reported following wounding in adult skin (Donati et al., 2017). However, it has not been technically possible to follow daughter cell fates long term following cell division with *ex vivo* imaging of embryonic explants.

Genetic lineage tracing is a powerful tool to identify clonally-related cells, and relies on induction of a permanent genetic mark—usually a fluorescent protein—by a tissue-specific, inducible Cre recombinase (Fig. 6A). Previously, we and others have used short-term “mitotic” genetic lineage tracing to characterize self-renewing and differentiating behaviors (Byrd et al., 2019; Poulson & Lechler, 2010; Williams et al., 2014). These include the following clone types: 1) symmetric cell divisions (SCD), which contain two basal cells; 2) asymmetric cell divisions (ACD), which contain one basal and one suprabasal cells; and 3) delaminations, which contain a single suprabasal (spinous) cell. Across these studies, there has been a good correlation between the ratio of SCDs to ACDS in clonal data with observations of division orientation in fixed tissue. However, whether loss of essential spindle orientation genes affects cell fate choices has never been directly shown.

**Figure 6.**
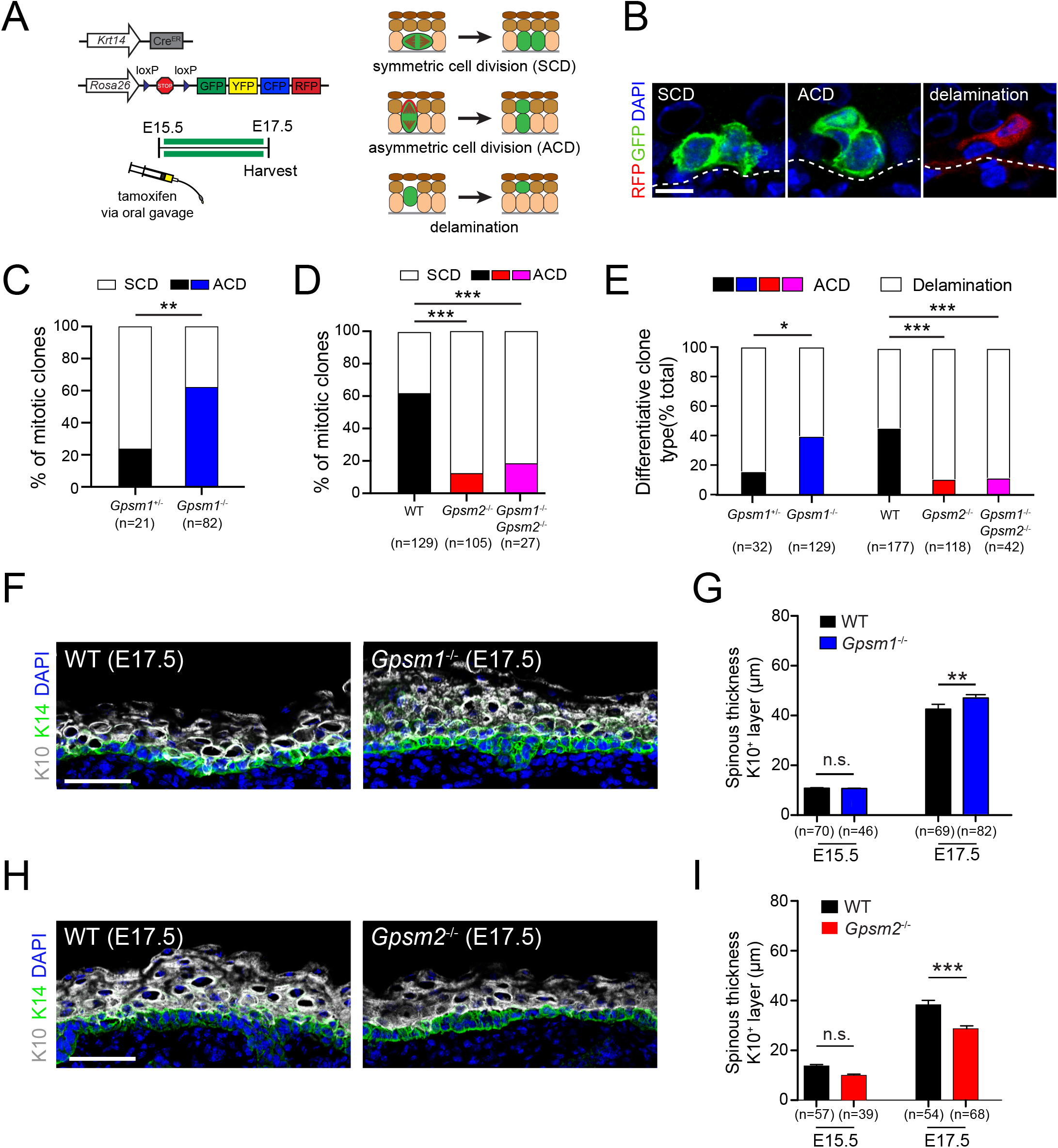
AGS3/Gpsm1 loss promotes asymmetric cell fates and differentiation. (A) Graphical depiction of clonal lineage tracing strategy using *Krt14*^*CreER*^ and the multi-colored *Rosa26*^*confetti*^ reporter, for data shown in (B-E). (B) Representative images of showing three types of clones. (C) Quantification of proportion of symmetric cell division (SCD) vs asymmetric cell division (ACD) clones in AGS3 KOs (*Gpsm1*^*-/-*^) (blue) compared to AGS3 heterozygote (*Gpsm1*^*+/-*^) controls (black). (D) Same analysis as (C) but for WT (black), LGN KO (*Gpsm2*^*-/-*^, red), and AGS3/LGN double KO (*Gpsm1*^*-/-*^; *Gpsm2*^*-/-*^, magenta) clones. (E) Proportions of differentiative clones occurring by ACD versus delamination for indicated genotypes. (F-I) Immunofluorescent images (F,H) and quantification (G,I) of spinous (K10, gray) layer thickness in AGS3 KOs (F,G) and LGN KOs (H,I) compared to WT littermate controls. Scale bars: 10µm in (B), 50 µm in (F,H); n values indicate clones (C-E) or measurements (G,I) from >3 independent embryos per genotype; n.s. = not significant, * p<0.05, ** p<0.01, *** p<0.001 by chi-square (C,D) or unpaired t-test (G,I).

Recently, we used mitotic genetic lineage tracing to show that loss of the adherens junction protein afadin (Afdn), which interferes with telophase reorientation, leads to an increase in asymmetric fate clones (Lough et al., 2019). Here, we apply a similar approach using *Krt14*^*CreERT2*^; *Rosa26*^*confetti*^ mice on LGN, AGS3 and double KO backgrounds, treated with a single low dose of tamoxifen at E15.5 and collected at E17.5 (Fig. 6A,B). Although the *Krt14* promoter is less active in suprabasal cells, in order to minimize inclusion of clones that may have been induced while already differentiated, single cell clones located above the first spinous layer were not counted as delamination events.

We compared the proportion of SCD and ACD clones in AGS3 KOs (*Gpsm1*^*-/-*^) and heterozygote (*Gpsm1*^*+/-*^) littermate controls, and observed a significant shift toward ACDs in KOs (Fig. 6C). On the other hand, loss of LGN led to a sharp increase in SCD clones (Fig. 6D). In further support of *Gpsm2* being epistatic to *Gpsm1*, AGS3/LGN double KOs showed a similar increase in SCD clones as LGN KOs (Fig. 6D). These data confirm that LGN and AGS3 play opposing roles in regulating oriented cell divisions, which result in altered cell fate choices at the clonal level.

It is worth noting that in both AGS3 and LGN mutants, there was excellent agreement between cell fate choices determined by genetic lineage tracing and division orientation at the “anaphase +1h” timepoint of our *ex vivo* live-imaging studies. For example, 61% of AGS3 KO cells showed perpendicular orientations at +1h (Fig. 4I), and 62% of divisions resulted in ACD clones (Fig. 6C). Similarly, in LGN KOs, 86% of imaged mitoses adopted planar orientations at +1h (Fig. 4F), and 88% of mitotic clones were SCDs (Fig. 6D). This suggests that *ex vivo* live imaging captures the behaviors that occur *in vivo*, and that measures of telophase orientations accurately reflect fate outcomes.

During epidermal development, differentiation can be accomplished through either ACD or delamination. Delamination is a process by which basal cells initiate differentiation within the basal layer (e.g., via upregulation of spinous keratins such as K10), followed by detachment from the underlying basement membrane and upward migration into the spinous layer (Ellis et al., 2019; Watt & Green, 1982; Wickstrom & Niessen, 2018; Williams et al., 2014). We and others have documented that delamination is the predominant mode of differentiation during early stratification (E12.5-E15.5), while ACDs become more common during peak to late stratification (Damen et al., 2021; Williams et al., 2014). In both embryonic and adult epidermis, the processes of proliferation and differentiation are spatially correlated and can locally influence each other (Mesa et al., 2018; Wickstrom & Niessen, 2018). However, it remains an open question whether mutations that alter the balance between SCDs and ACDs could impact delamination.

To test this, we examined whether the proportions of the two types of “differentiative” clones—ACD and delamination—were affected by LGN or AGS3 loss. Differentiation by delamination was significantly decreased in AGS3 KOs, which show a higher proportion of ACDs (Fig. 6E). Conversely, delamination frequency was significantly elevated in LGN KOs and AGS3/LGN double KOs, where ACDs are infrequent. Of note, the higher proportion of delamination clones in the AGS3 cohort is consistent with these mice being at a younger developmental stage compared to the LGN cohort, which could also explain the higher proportion of SCDs observed in *Gpsm1*^*+/-*^controls compared to *Gpsm2*^*+/+*^ controls (compare black bars in Fig 6C vs 6D). Nonetheless, these data demonstrate an inverse correlation between ACD prevalence and delamination, suggesting that these two differentiation mechanisms may be coordinated across the epithelium.

Finally, we examined global stratification in AGS3 KOs, LGN KOs and littermate controls using the differentiation marker K10 (Fig. 6F-I). No changes in spinous thickness were observed in either mutant at E14.5, consistent with the lack of any effect on spindle orientation by either gene at this age. However, by E16.5 we observed a significant decrease in spinous thickness in LGN KOs compared to WT littermates, consistent with previous observations in *Gpsm2*^*1617*^ knockdowns (Williams et al., 2011). While the effect was milder in AGS3 KOs, we also observed a significant increase in spinous thickness at E16.5 (Fig. 6G). Thus, during epidermal development, AGS3 promotes planar divisions, and its loss leads to altered fate choices and increased differentiation.

## DISCUSSION

Collectively, these studies show that LGN/Gpsm2 and AGS3/Gpsm1 play opposing roles in regulating oriented cell divisions and fate choices in the developing epidermis (Figure 7). Static and live analyses of division orientation, genetic lineage tracing and quantification of differentiation markers confirm that AGS3 promotes self-renewal through SCDs while LGN promotes differentiation via ACDs. LGN knockdown (Williams et al., 2011) or knockout (these studies) leads to a planar bias in division orientation, which we also observe when AGS3 is overexpressed. Conversely, AGS3 loss has the reverse effect, increasing perpendicular divisions. LGN, Pins and its homologs have evolutionarily-conserved roles in determining the axis of the mitotic spindle. Beyond this canonical function, we now provide evidence that LGN is also required during late stages of mitosis to correct “errant” oblique divisions toward perpendicular. Interestingly, in the absence of AGS3 we do not observe any obvious defects in the initial apical positioning of LGN, nor are division angles significantly altered at anaphase onset. Thus, the strongest effects of AGS3 in promoting planar divisions appear to occur during telophase correction. Future studies will be necessary to determine why and how AGS3 functions during this specific phase of the cell cycle.

**Figure 7.**
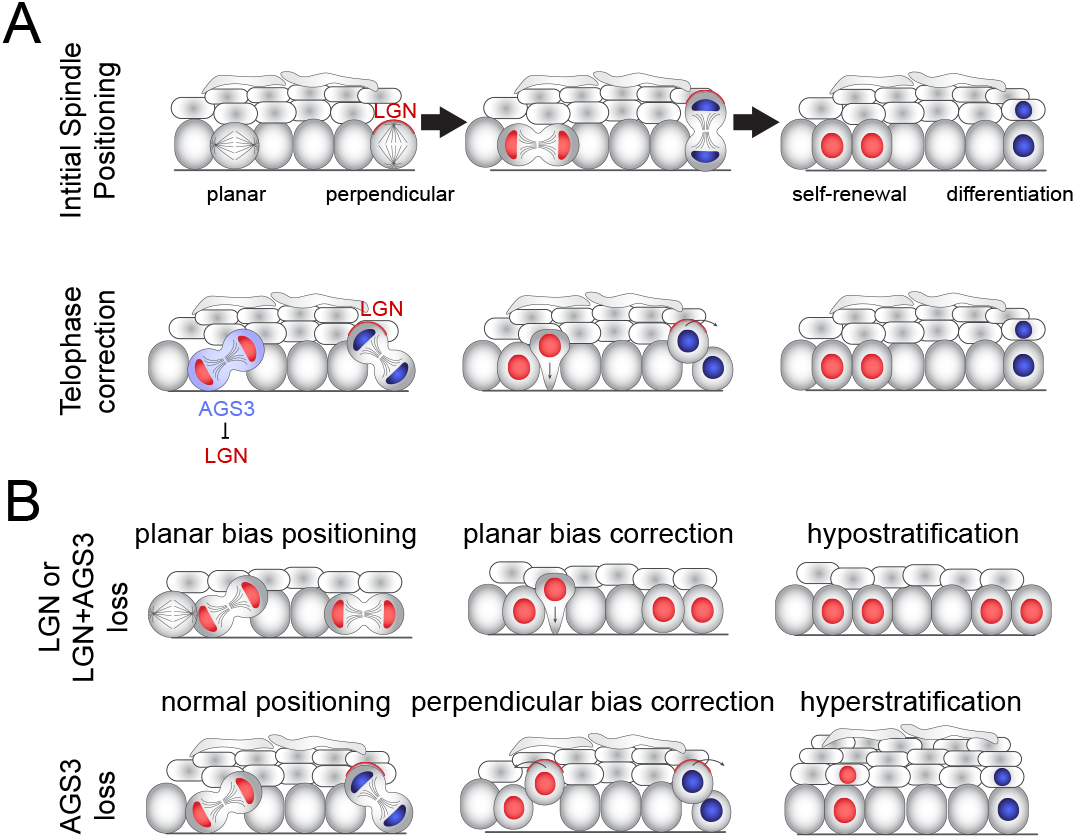
Model of two-step process for oriented cell divisions. (A) Basal progenitors can undergo either self-renewing planar symmetric divisions (red cell) or differentiative perpendicular asymmetric divisions (blue cell). Oriented cell divisions are regulated in a two-step process: 1) initial spindle positioning (top) and 2) telophase correction (bottom). Polarized apical LGN promotes apical spindles at metaphase and also promotes perpendicular reorientation during telophase. Primarily cytoplasmic AGS3 displaces LGN from the cortex to promote planar correction. (B) Phenotypes resulting from loss of LGN and/or AGS3.

### Mechanisms of competition between LGN and AGS3

In theory AGS3 could promote planar divisions through a variety of mechanisms, to be discussed in greater detail below. However, we provide the following lines of evidence that AGS3 acts as a novel negative regulator of LGN: 1) AGS3 is enriched in the cytoplasm, and thus is unlikely to interact directly with the cortical spindle orientation machinery, 2) AGS3 loss- or gain-of-function promotes or inhibits, respectively, the apical localization of LGN, 3) *Gpsm2* is epistatic to *Gpsm1*, demonstrating that LGN loss masks AGS3 function, and implying that AGS3 acts through LGN.

In thinking about how GPSM complexes regulate spindle orientation, it is important to consider their accessibility, partner binding affinities, and stoichiometry of their interactions. First, free GPSM proteins exist in a closed confirmation, where the N-terminal TPR region is bound to the C-terminal GPR repeats. The only protein known to be capable of binding LGN in its closed conformation is Gαi—preferentially in its GDP-loaded form—which is able to access the first GoLoco motif that is not involved in intramolecular interactions (Du & Macara, 2004; Nipper, Siller, Smith, Doe, & Prehoda, 2007; Takayanagi et al., 2019). Thus, Gαi-GDP binding is thought to be an early event in complex formation, promoting cortical association of GPSM proteins due to Gαi myristoylation, and catalyzing additional Gαi binding that promotes the open conformation. An important corollary of the Gαi-GDP preference of GPR domains is that GPSM proteins act as guanine dissociation inhibitors that reduce the exchange of GDP for GTP on Gαi subunits, which in some contexts can be opposed by the guanine nucleotide exchange factor Ric-8A (David et al., 2005; Hampoelz, Hoeller, Bowman, Dunican, & Knoblich, 2005; Woodard et al., 2010). Second, structural studies have shown that LGN TPR domains cannot simultaneously bind multiple proteins, e.g. NuMA and Insc (Culurgioni et al., 2011; Yuzawa et al., 2011; Zhu, Wen, et al., 2011). The current model is that Insc helps recruit LGN to the apical cortex but LGN later becomes released from Insc—by mechanisms that are still not well understood—to enable it to bind to NuMA (Culurgioni & Mapelli, 2013). Third, NuMA can bind both microtubules and dynein directly and is thought to be anchored by the LGN-NuMA-Gαi ternary complex at the plasma membrane to ensure correct spindle positioning (Kotak, Busso, & Gonczy, 2012; Seldin et al., 2016). Finally, structural studies have shown that LGN and NuMA can exist in a heterohexameric ring structure, which forms higher order networks that facilitate microtubule capture and orient spindles in mammalian cells (Pirovano et al., 2019).

At a molecular level, we favor a model in which AGS3 competes with LGN for NuMA binding, preventing LGN from forming productive ternary complexes that capture astral microtubules at the apical cortex. This hypothesis is based on several lines of evidence. First, there is direct biochemical evidence that AGS3 can bind to NuMA (Saadaoui et al., 2017), and the specific residues within the TPR5/6 domains of LGN that mediate binding to NuMA—N203, R221 and R236 (Culurgioni et al., 2011; Zhu, Wen, et al., 2011)—are conserved in AGS3. Second, a construct consisting of the AGS3-TPR domains localizes to spindle poles—where NuMA is present— while the TPR domains of LGN localize to the cytoplasm (Saadaoui et al., 2017). Moreover, an LGN chimera containing the AGS3 TPR domain can displace WT LGN from the cell cortex but can only partially rescue a Gpsm2 spindle orientation defect (Saadaoui et al., 2017). This suggests that the TPR domains of AGS3 might have a higher affinity for NuMA than LGN, but cannot orient spindles efficiently, perhaps because AGS3-NuMA cannot form a functional apical complex. Third, mutations in LGN, or NuMA, that render them oligomerization deficient—e.g., unable to form hexameric ring structures—lack spindle orienting ability (Pirovano et al., 2019). Thus, the ability of GPSM protein TPR domains to form higher order complexes with NuMA appears to be critical for their function.

These studies also showed that the curvature of the NuMA-LGN hexamer is accomplished by the unusually long fourth TPR domain of LGN—which contains 54 amino acids instead of 34 (Pirovano et al., 2019). Interestingly, murine AGS3 shows the poorest conservation with LGN in this particular domain (63% similar compared to >86% similarity for all other TPRs), with the highest dissimilarity is in the region between the two alpha-helical regions (data not shown). While the crystal structure of the AGS3 TPR has not been solved, the divergence of this particular TPR repeat that is critical for ring formation suggests that AGS3 may not be capable of forming heterohexamers with NuMA. Moreover, another region of LGN that is critical for its oligomerization capacity is the short N-terminus that precedes the first TPR repeat (Pirovano et al., 2019). This N-terminal region is highly conserved among vertebrate Gpsm2 orthologs, but entirely divergent among Gpsm1 orthologs (data not shown), providing a second line of evidence that NuMA bound to AGS3 is unlikely to form ring structures.

While we favor that LGN and AGS3 compete for NuMA binding, it is also possible that AGS3 could bind to, and inhibit, LGN directly. For example, it has been shown that the AGS3 GPR region can pull-down the TPR domain of LGN (Saadaoui et al., 2017). Unfortunately, crossreactivity between our LGN and AGS3 antibodies makes it difficult to test for colocalization in mitotic basal cells, as even our reasonably specific guinea pig anti-LGN antibody appears to label some AGS3 in LGN KOs (Fig. 1A). Nonetheless, because AGS3 is primarily cytoplasmic, it could sequester a pool of LGN away from the cell cortex, preventing engagement with NuMA. One caveat to this model is that in order for AGS3 to bind to LGN, they would both need to be in their open conformations, a process believed to require Gαi-GDP binding. Because Gαi proteins are myristoylated and membrane-associated, this event would be more likely to occur at the cell cortex.

### Binding partners and subcellular localization of LGN and AGS3

The spindle orienting function of *Gpsm2* and its homologs is highly conserved throughout evolution, and may have evolved as early as the time when bilateria and cnidaria diverged (Schiller & Bergstralh, 2021; Wavreil & Yajima, 2020). While LGN and its orthologs have an evolutionarily-conserved role in oriented cell divisions, the manner and pattern in which LGN localizes to the cell cortex varies among epithelial tissues. For example, in the retina and developing oral epithelia, LGN localizes apically and serves a similar function in promoting perpendicular divisions as in epidermis (Byrd et al., 2016; Lacomme et al., 2016). On the other hand, in the tactile filiform papilla of the dorsal tongue, radial glia of the subventricular zone, and neural tube, LGN localizes laterally and promotes planar divisions (Byrd et al., 2016; Konno et al., 2008; Lacomme et al., 2016; Morin et al., 2007; Peyre et al., 2011). Thus, while the spindle orienting capacity of LGN is conserved, its variable subcellular localization dictates division directionality differently across tissues.

In addition to Insc and Gαi family proteins, a growing list of proteins— including Dlg, E-cadherin, Frmpd1/4 and afadin—have been found to promote the cortical localization of LGN in different contexts (Carminati et al., 2016; Gloerich, Bianchini, Siemers, Cohen, & Nelson, 2017; Schiller & Bergstralh, 2021; Wee, Johnston, Prehoda, & Doe, 2011; Yuzawa et al., 2011). Although autoinhibition and phosphorylation are mechanisms known to regulate LGN’s ability to interact with proteins that promote its membrane association (Bergstralh et al., 2017; Johnston, Hirono, Prehoda, & Doe, 2009; Pan et al., 2013), comparatively less is known about negative regulators. One example is SAPCD2, which inhibits the cortical localization of LGN in the retina, possibly by competing for NuMA binding (Chiu et al., 2016). We believe that AGS3 is another, though clearly its function is context-dependent, given its inability to orient spindles or influence LGN localization in neurons (Saadaoui et al., 2017).

While unbiased proteomic approaches might best ascertain how complexes containing AGS3 and LGN differ, we speculate that differences in their subcellular localization and function could also be attributable to posttranslational modifications such as phosphorylation. The flexible linker domain of LGN/Pins has been shown to be phosphorylated by both aPKC and Aurora kinase, which mediate interactions with 14-3-3 and discs large (Dlg), respectively (Hao et al., 2010; Johnston et al., 2009). While it is not known whether aPKC phosphorylation of LGN impacts its spindle orienting capacity, loss of *Prkci* in the epidermis does result in a spindle orientation phenotype (Niessen et al., 2013). On the other hand, the Dlg-LGN interaction is highly conserved throughout evolution (Schiller & Bergstralh, 2021), and a non-phosphorylatable *pinsS436A* mutant is non-functional in a *Drosophila* S2 induced-polarity assay, while the equivalent mutation in vertebrate LGN (S401A) induces spindle misorientation in MDCK cells and the chick neural tube (Hao et al., 2010; Saadaoui et al., 2014). This serine residue is conserved in the AGS3 linker, but there are some differences in the flanking residues, and it has been reported that the AGS3 linker binds Dlg with ∼500-fold lower efficiency than LGN (Zhu, Shang, et al., 2011). While substitution of the LGN linker into AGS3 is not sufficient to relocalize AGS3 from the cytoplasm to the cortex (Saadaoui et al., 2017), the converse experiment has not been attempted. Other potential phosphorylation sites for LGN include T450, which promotes growth in breast cancer cells (Fukukawa, Ueda, Nishidate, Katagiri, & Nakamura, 2010).

Another way in which AGS3 and LGN might differ is through their interactions with G-proteins. Although LGN and AGS3 bind Gαi proteins with similar affinity in vitro (Willard et al., 2008), some of the best insights into how Gαi interactions influence their localization and function *in vivo* come from domain swap experiments between the GPR regions of LGN and AGS3 (Saadaoui et al., 2017). The Morin lab found that while the AGS3 GPR domain cannot substitute for the LGN GPR domain, replacing the inter-GPR regions of AGS3 with those of LGN confers cortical localization. The converse is also true, in that replacing the inter-GPR regions of LGN with AGS3’s results in cytoplasmic localization. Moreover, for AGS3 it has been shown that addition of inter-GPR regions significantly enhances Gαi1 binding compared to isolated GPR domains alone (Adhikari & Sprang, 2003). Collectively, these findings suggest that the inter-domain regions contain important information that might regulate the ability of LGN/AGS3 to interact with specific Gαi proteins.

Saadaoui and colleagues speculated that variability in the interdomain regions might differentially affect intramolecular interactions between the TPR and GPR domains of LGN and AGS3, but conceivably, this could also impact their ability to bind specific Gαi proteins, particularly if post-translational modifications occur within these interdomains. There are three Gαi proteins in mammals and all are expressed in the epidermis, but it is Gαi3 which colocalizes with LGN, and Gnai3 loss leads to a planar bias like loss of LGN (Williams et al., 2014). Gnai2 loss has no obvious phenotype (Williams et al., 2014), and Gnai1 loss has yet to be explored, so it remains possible that AGS3 may preferentially associate with a different cohort of Gαi proteins than LGN.

### Are planar divisions an active or “default” process?

While much remains to be explored at the molecular level of how AGS3 interacts with the spindle orientation machinery, our studies shed new light on both the negative and positive regulation of perpendicular, asymmetric divisions. Importantly, we view AGS3 as a negative regulator of perpendicular divisions, rather than a promoter of planar divisions. We speculate that relative levels of AGS3 and LGN within individual cells determine the likelihood that they will both establish and maintain a perpendicular orientation throughout mitosis. However, this cannot explain why a significant fraction of basal cells “choose” planar orientations independent of AGS3. The fact that LGN loss—and also combined AGS3/LGN loss—results in a strong planar bias rather than randomization of division angles suggests that either 1) there is an as-yet undiscovered active mechanism to promote planar divisions, or 2) planar may be the default or passive process.

We favor the latter hypothesis for the following reasons. First, LGN is necessary for NuMA’s cortical localization, and NuMA—or dynactin— loss leads to a similar phenotype as LGN loss in the epidermis (Seldin et al., 2016; Seldin, Poulson, Foote, & Lechler, 2013; Williams et al., 2011). While there are numerous examples in the literature where NuMA can direct spindle orientation in an LGN/Pins-independent manner (Bergstralh et al., 2016; Bosveld et al., 2016; Kiyomitsu & Cheeseman, 2013; Kotak, Busso, & Gonczy, 2014), there are few examples where spindles can be actively reoriented independently of NuMA and dynein. Second, planar divisions predominate in the early epidermis, prior to and during early stratification, and we are not familiar with any genetic alteration that induces precocious perpendicular divisions or stratification (not to be confused with the hyperstratification that is observed in *Gpsm1* and *Afdn* mutants).

More likely, we believe that planar divisions are a default state, attributable to high tension across the basal layer, possibly as a result of stronger adhesive forces on lateral versus apical cell membranes or intra-tissue tension provided by differentiated layers (Ning, Muroyama, Li, & Lechler, 2021). Of interest, Devenport and colleagues recently described increased perpendicular divisions and hyperthickened epidermis in *Vangl2* mutants (Box, Joyce, & Devenport, 2019). Unexpectedly, this phenotype was not due to defective planar cell polarity, but rather to tissue-wide changes in cell shape and packing caused by the neural tube defect. In addition to this effect of interphase cell shape on division orientation, hypoproliferation and elevated apoptosis can lead to non-cell autonomous increases in planar divisions (Morrow, Underwood, Seldin, Hinnant, & Lechler, 2019; Soffer et al, under revision). Future studies will be necessary to directly test how local changes in cell density, and intra-tissue tension, impact oriented cell divisions and the balance between proliferation and differentiation.

### Coordinating self-renewal and differentiation on a tissue scale

In the epidermis, ACDs are one of two known mechanisms to promote differentiation, the other being delamination (Watt & Green, 1982). Until 15 years ago, delamination—e.g. differentiation by detachment rather than division—was thought to be the driving force for epidermal differentiation (Blanpain & Fuchs, 2006). While this is true during adult epidermal homeostasis, where perpendicular divisions are rare to non-existent (Clayton et al., 2007; Ipponjima, Hibi, & Nemoto, 2016; Mesa et al., 2018; Rompolas et al., 2016), it is now clear that ACDs are essential for proper skin development in the embryo (Lechler & Fuchs, 2005; Seldin et al., 2016; Williams et al., 2014). In the adult, the processes of proliferation and differentiation cooperate to regulate cell density in the basal layer, such that delamination precedes and induces self-renewal in nearby cells (Cockburn et al., 2021; Mesa et al., 2018). Interestingly, in the embryo, the opposite relationship exists, such that proliferation seems to drive neighboring cells to delaminate (Miroshnikova et al., 2018). In other tissues, local crowding drives live-cell extrusion (Eisenhoffer et al., 2012; Marinari et al., 2012), and “winners” (stem cells) and “losers” (differentiating cells) have also been described during epidermal stratification (Ellis et al., 2019). Yet, whether the two differentiation processes of ACD and delamination are linked has not been explored. Here, using genetic lineage tracing, we find that delamination frequency decreases when ACDs increase in AGS3 KOs, and conversely, delamination increases when ACDs decrease in LGN KOs. It will be of interest to see how these global changes are influenced by the local tissue microenvironment.

## MATERIALS AND METHODS

### Animals

All mice were housed in an AAALAC-accredited (#329; November, 2020), USDA registered (55-R-0004), NIH welfare-assured (D16-00256 (A3410-01) animal facility. All procedures were performed under IACUC-approved animal protocols (19-155 and 22-121). For fixed sample imaging (immunohistochemistry) and all lentiviral transduction experiments (unless otherwise noted), CD1 wild-type outbred mice (Charles Rivers; #022) were utilized. *Gpsm2-/-*KO (*LGN*^*BF*^ (*Gpsm2tm1a*(EUCOMM)Wtsi; Jackson Labs #4441912 via Basile Tarchini) (Tarchini et al., 2013) mice were maintained on a mixed C57B6/J background and bred to either 1) *Krt14*^*CreER*^; *Rosa26*^*Confetti*^ (*Tg(KRT14-cre/ERT)20Efu*; Jackson Labs #005107 / *Gt(ROSA)26Sortm1(CAG-Brainbow2*.*1*)Cle; Jackson Labs #013731) females or identical males for genetic lineage tracing, and 2) *Rosa26*^*mTmG*^ (*Gt(ROSA)26Sortm4(ACTB-tdTomato,-EGFP*)Luo/J; Jackson Labs #007576) homozygous females with at least one copy of the *Krt14*^*Cre*^ allele (crossed to males of the identical genotype), for *ex vivo* live imaging. *Gpsm1-/-*KO mice (*AGS3DEL(B6*.*129S6(SJL*)-*Gpsm1tm1*.*1Lajb/J*; Jackson Labs # 019503 via Ricardo Richardson) (Blumer et al., 2008) were maintained on a mixed 129S6 background and bred to the same strains as *Gpsm2* KOs for lineage tracing and live-imaging. *Gpsm1-/-; Gpsm2-/-*mice were maintained on a mixed background and bred to the same strains as *Gpsm2* KOs for lineage tracing and live-imaging. CD1, *Gpsm2* KO, and *Rosa26*^*mT/mG*^; *Krt14*^*Cre*^ animals were injected with lentiviral constructs (see below). Note that all ages are defined where E0.5 is noon of the day that a plug is found, but that developmental differences exist between strains. For example, we have found the 129S6 background of AGS3 KOs to be ∼1d behind, and the C57Bl6/J background of LGN KOs to be ∼0.5d behind outbred CD1s.

### Live Imaging

The protocol for live imaging has been adapted from the technique described by the Devenport lab (Cetera et al., 2018). For a full protocol please see (Lough et al., 2019). Briefly, epidermal explants were harvested from the mid-back of wild-type E16.5 *Rosa26*^*mT/mG*^; *Krt14*^*Cre*^ embryos, crossed to either *Gpsm2* KO, *Gpsm1* KO, or double mutant background. The explants were sandwiched between a gel/media patty and a gas-permeable membrane dish, and were cultured at 37°C with 5.0% CO2 for >1.5 hours prior to- and throughout the course of imaging. Confocal imaging was used to acquire a 20-30 micron z-stack every 5 min for 3-6h in a temperature controlled chamber, with the exception of images in Fig. 2E,F, which were acquired using the Dragonfly spinning disk confocal (Andor) equipped with a Leica 40X/1.4 NA Oil Plan Neo objective. Images were acquired with 5 minute intervals and a Z-series with 0.5 mm step-size (total depth ranging from 46 microns) for 6 hours. The Andor iXon life 888 BV was used to image mKate2, Andor Zyla 4.2 Plus was used for imaging mGFP. Additionally, we performed live imaging of E16.5 lentiviral-transduced *Gpsm1*^*1147*^ H2B-mRFP1 epidermal explants on a *Rosa26*^*mT/mG*^; *Krt14*^*Cre*^ background. Divisions appearing close to the tissue edge or showing any signs of disorganization/damage were avoided to exclude morphological changes associated with wound-repair. 4D image sets were processed with a deep-learning (DL) image denoising method using self-supervised training called Noise2Void (detailed bellow), and processed using ImageJ (Fiji).

### Lentiviral Injections

The protocol for lentiviral injection was performed according to (Beronja et al., 2010), under approved IACUC protocol 19-155. Pregnant mice mice were anesthetized for less than 1h and provided subcutaneous analgesics (5 mg/kg meloxicam and 1-4 mg/kg bupivacaine). A uterine horn was pulled out the mom into a PBS filled culture dish to expose E9.5 embryos. We performed a microinjection of ∼0.7 µl of concentrated lentivirus into the amniotic space using a custom glass needle that was visualized by ultrasound. Three to six embryos were injected on the same horn per pregnant dam, and the non-injected horn was used for matched littermate controls. Following injection, the horns were put back into the thoracic cavity of the dam and sutured closed. Surgical staples were used to reseal the skin incision. Once awake and freely moving, the dam was monitored for 4-7 days. Embryos were harvested and processed at E14.5-E17.5.

### Genetic Lineage Tracing

For full protocol, please see (Lough et al., 2019). Males of identical genotype were crossed with *Krt14*^*CreERT2*^; *Rosa26*^*Confetti*^ females with either *Gpsm2* KO, *Gpsm1* KO or double mutant genotypes. A 100 µg per gram dam mass of tamoxifen was delivered by oral gavage at E15.5 to activate the *Krt14*^*CreERT2*^ allele. Following tamoxifen dosing dams were monitored for 24h for signs of abortion or distress. 48h after tamoxifen delivery, embryos were harvested at E17.5, backskins were fixed for 30 min in 4% PFA and washed with PBS. Fixed backskins were embedded in OCT and sectioned sagittally at 8 µm. To enhance the fluorophores of the *Rosa26*^*Confetti*^ allele we immunostained backskin sections (see below), omitting the 5 min post-fixation step, using monoclonal Rat anti-mCherry clone 16D7 (Life Technologies M11217) to enhance membrane-RFP, and polyclonal Chicken anti-GFP (Abcam ab13970) to enhance the membrane-CFP, nuclear-GFP, and cytoplasmic-YFP fluorophores. Areas of the stained section with labeled clones were acquired with a 40x/1.15 NA objective with 1.5x digital zoom. We scored sparse clones (<1% total cells) for the number of suprabasal cells (distinguished by staining with anti-Krt10 antibody), and basal cells (distinguished by staining with anti-Krt14 antibody). To exclude the possibility that tamoxifen induction occurred while cells were already suprabasally positioned, we only counted delamination events as clones with suprabasal cells in the stratum spinosum (SS) layer.

### Constructs and RNAi

For *Gpsm2* and *Gpsm1* RNAi targeting, we utilized shRNAs that had been previously validated (Williams et al., 2011; Williams et al., 2014). The nucleotide base (NCBI Accession number) for a given shRNA clone are identified by the gene name followed by the 21-nucleotide target sequence (e.g. *Gpsm1*^*1147*^). Packaging of lentivirus was performed using 293FT cells and pMD2.G and psPAX2 helper plasmids (Addgene plasmids #12259 and #12260, respectively). To evaluate shRNAs for their knockdown efficiency, we infected primary keratinocytes with a MOI of ∼1 in E-Low calcium medium for ∼48 h. Puromycin selected cell lines were lysed with RNeasy Mini Kit (Qiagen) to isolate RNA. mRNA knockdown efficiency was determined by RT-qPCR using 2 independent primer sets for each transcript with *Hprt1* and cyclophilin B (*Ppib2*) as reference genes and cDNA from stable cell lines expressing Scramble shRNA as a reference control. The following primer sequences were used: *Scramble* (5’-CAACAAGATGAA-GAGCACCAA-3’), *Gpsm1*^*1147*^ (5’-GCCTTGACCTTTGCCAAGAAA-3’). Sequences for *Hprt1* and *Ppib2* primers have been described previously (Williams et al., 2011).

### Antibodies, Immunohistochemistry and Fixed Imaging

E15.5-E16.5 embryos were skinned and flat-mounted on Whatman paper. E14.5 embryos were mounted whole. In all cases (except for lineage tracing experiments), samples were mounted in OCT (Tissue Tek) and frozen fresh at -20°C. Experimental genotypes (homozygous, heterozygous and wild-type) were mounted in the same block to allow for direct comparisons on the same slide. Similarly, infected and uninfected littermate controls were mounted in the same blocks. Frozen samples were sectioned (8 µm thickness) on a Leica CM1950 cryostat. Staining was conducted as previously described (Lough et al., 2019). Images were acquired using LAS AF software on a Leica TCS SPE-II 4 laser confocal system on a DM5500 microscope with a ACS Apochromat 40x/1.15 NA oil, or ACS Apochromat 63x/1.30 NA oil objectives. The following primary antibodies were used: Survivin (rabbit, Cell Signaling 2808S, 1:500), LGN (guinea pig, 1:500) (Williams et al., 2011), LGN (rabbit, Millipore ABT174, 1:2000) (Williams et al., 2014), phospho-histone H3 (rat, Abcam AB10543, 1:1000-5000), β4-integrin (rat, ThermoFisher 553745, 1:1,000), Cytokeratin-14 (chicken, Biolegend 906004, 1:5000), Cytokeratin-14 (guinea pig, Origen BP5009, 1:1000), Cytokeratin-10 (rabbit, Covance 905401, 1:1000), GFP (chicken, Abcam AB13970, 1:1000), V5 (chicken, Abcam AB9113, 1:1000), mCherry (rat, Life Technologies M11217, 1:1000-3000), RFP (rabbit, MBL PM005, 1:1000), RFP (chicken, Millipore AB3528, 1:500). The following secondary antibodies were used (all antibodies produced in donkey): anti-rabbit AlexaFluor 488 (Life Technologies, 1:1000), anti-rabbit Rhodamine Red-X (Jackson Labs, 1:500), anti-rabbit Cy5 (Jackson Labs, 1:400), anti-rat AlexaFluor 488 (Life Technologies, 1:1000), anti-rat Rhodamine Red-X (Jackson Labs, 1:500), anti-rat Cy5 (Jackson Labs, 1:400), anti-guinea pig AlexaFluor 488 (Life Technologies, 1:1000), anti-guinea pig Rhodamine Red-X (Jackson Labs, 1:500), anti-guinea pig Cy5 (Jackson Labs, 1:400), anti-goat AlexaFluor 488 (Life Technologies, 1:1000), anti-goat Cy5 (Jackson Labs, 1:400), anti-mouse IgG AlexaFluor 488 (Life Technologies, 1:1000), anti-mouse IgG Cy5 (Jackson Labs, 1:400), anti-mouse IgM Cy3 (Jackson Labs, 1:500).

### Image Processing, Measurements, Quantification, and Statistics

Denoising: Live imaging acquisition on epidermal explants was conducted on a Zeiss LSM 710 Spectral confocal laser scanning microscope as detailed in (Lough et al., 2019). To restore noisy images we employed Noise2Void (N2V), a deep-learning (DL) image denoising method using self-supervised training (Krull et al., 2020). While this method does not require ground truth or low noise equivalent images to train a DL model, we nevertheless validated our N2V models by error mapping and quality metrics estimation of ground truth images (SSIM and RSE maps).

### Spindle and Division Orientation

Spindle orientation measurements was performed as in (Byrd et al., 2016; Lough et al., 2019; Williams et al., 2011; Williams et al., 2014). Cells in metaphase were identified based on nuclear morphology. Anaphase cells were determined by both nuclear condensation and broadly distributed Survivin staining between daughter cells. Telophase cells were identified due to dual-punctate Survivin staining. Division orientation measurements for anaphase and telophase were calculated as the angle between a vector parallel to the basement membrane and a vector connecting the estimated center of each daughter nuclei. Similarly, division orientation was measured in live imaging experiments, where the position of the daughter nuclei was inferred based on cell volume/shape changes.

### LGN/AGS3 localization/intensity

Imaging of CD1 lentiviral H2B-RFP animals was performed with control littermates and experimental samples on the same slide to avoid variation in antibody staining. Similarly, with experimental genotypes, imaging was performed with control littermates on the same slide. Scoring of LGN localization patterns (e.g. apical, unpolarized, absent) was determined for cells labeled with pHH3 or Survivin (telophase marker), irrespective of the lentiviral H2B-RFP reporter to avoid bias. Whole cell LGN/AGS3 levels were measured from a single z-slice determined to be the cell center. The cell edge was inferred based on K14 signal exclusion, and plotted as integrated intensity.

### Stratification

Epidermal thickness was measure in E14.5-E17.5 embryos by staining for H2B-RFP, K14 and K10. We imaged >10 regions of sagittal backskin sections per quantified embryo. We thresholded the K10 signal and created a binary mask that was used to quantify the suprabasal area above threshold. The area was then divided by the length of the underlying basement membrane (determined by K14 staining). n values for these analyses are representative of the number of regions imaged.

### Statistical analyses and graphs

Error bars represent standard error of the mean (s.e.m.) unless otherwise noted. Statistical tests of significance were determined by Mann-Whitney U-test (non-parametric) or student’s t-test (parametric) depending on whether the data fit a standard distribution (determined by pass/fail of majority of the following: Anderson-Darling, D’Agostino & Pearson, Shapiro-Wilk, and Kolmogorov-Smirnov tests). Cumulative frequency distributions were evaluated for significant differences by Kolmogorov-Smirnov test. All analyses were generated using GraphPad Prism 9. Figures were constructed using Fiji and Adobe Illustrator CC 2021.

## Acknowledgements

We thank Basile Tarchini (Jackson Laboratories) for sharing the *Gpsm2* mice and Ricardo Richardson (North Carolina Central University) and Steve Lanier (Wayne State University) for sharing the *Gpsm1* mice. We are grateful to Ali Vural (Wayne State University) for sharing AGS3 antibodies and for helpful discussions. We thank Juliet King, Bethany Brown and Kevin Byrd for helpful discussions and technical support, as well as Bob Goldstein, Joan Taylor, Jim Bear and Stephanie Gupton for critical reading of the manuscript. We thank Jabiz Nafisi and Carien Niessen for their insights and sharing unpublished data. We are grateful to Pablo Ariel and the Microscopy Services Laboratory core facility for assistance and guidance with live imaging and image processing. These studies were supported by grants from the NIH R01 AR077591 (S.E.W), Binational Science Foundation 2019230 (S.E.W.), Sidney Kimmel Foundation Kimmel Scholars Program SKF-15-065 (S.E.W.), and Chan Zuckerberg Initiative DAF, an advised fund of Silicon Valley Community Foundation grant 2020-225716 (K.M.K). The MSL is supported in part by P30 CA016086 Cancer Center Core Support Grant to the UNC Lineberger Comprehensive Cancer Center, and the Andor Dragonfly microscope was funded with support from NIH grant S10 OD030223.

## Declaration of Interests

We have no interests or conflicts to declare.

## Notes

### Competing Interest Statement

The authors have declared no competing interest.

